# Characterizing population structure and documenting rapid loss of genetic diversity in Chiricahua Leopard Frogs (*Lithobates chiricahuensis*) with high throughput microsatellite genotyping

**DOI:** 10.1101/2024.05.31.596843

**Authors:** Caleb Beimfohr, Linet Rivas Moreno, Regan Anderson, Raina Cardwell, Zach Seeman, Ryan Spry, Matthew L Holding, Audrey Owens, Robert Denton

## Abstract

The use of molecular markers to assess genetic diversity has become a common component of recovery action plans for threatened and endangered species. In this study, we use an unusually large number of microsatellite markers (N=91) to characterize the genetic variation of Chiricahua Leopard Frogs (*Lithobates chiricahuensis*) across their range in order to understand their distribution of genetic variation, identify genetic bottlenecks, and measure genetic changes over time in a single, highly-managed population. Populations were best divided into three genetically distinct clusters, with the southeastern Arizona and New Mexico populations forming distinct genetic clusters. While there is moderate genetic variation distributed across the sampled populations, each population on its own shows relatively low allelic diversity. Most populations displayed strong genetic signals of recent genetic bottlenecks or a deficiency of heterozygous genotypes that is typically associated with frequent inbreeding. Populations that have a history of no management through translocations harbored the greatest number of unique alleles and overall allelic richness, especially in a subset of the Mexican populations. Finally, long-term cohort sampling at one specific site (the Southwestern Research Station in Portal, Arizona) allowed us to demonstrate how rapidly genetic diversity can decrease across a matter of years in a population with few founders. This work shows how microsatellite markers can provide important context for conservation agencies, but even a large suite of markers beyond what is typical may not be enough for populations that are extremely bottlenecked and have low levels of standing genetic diversity.

## Introduction

Low population genetic diversity can lead to a decrease in overall fitness and increase the risk of extinction (Soulé and Mills 1998; Reed and Frankham 2003). For this reason, estimating genetic diversity has been an important component of species conservation plans. Information on a species’ genomic variation and relatedness among populations can help inform conservation management decisions (Rivers et al. 2014). For certain scenarios, this may mean preserving the integrity of populations that contain unique genomic variation (Coates et al. 2018). For others, this information may guide reintroduction efforts or human-assisted dispersal (Whiteley et al. 2015; Jangjoo et al. 2016; Bell et al. 2019). Accurate population genetic measurements are at the center of strategies like genetic rescue, where immigrant individuals are purposely added to populations in an effort to restore heterozygosity and allelic richness (Hedrick et al. 2011). Successful examples include the Florida panther (*Puma concolor coryi;* Johnson et al. 2010), Swedish adder (*Vipera berus*; Madsen et al. 1999), and the greater prairie chicken (*Tymphanicus cupido*; Westemeier et al. 1998).

The techniques used to sample genomic variation have evolved rapidly in the past several decades (Mardis 2011). One step along this path—microsatellites—have outlasted many other approaches because they provide a large amount of information-per-locus (Kalinowski 2002) and allow for the re-use of primers that were initially large investments (Selkoe and Toonen 2006; Peterman et al. 2012). Although using genome-wide sequencing and single nucleotide polymorphisms (SNPs) can be more powerful and easier to scale, the availability of genomic resources across animals varies drastically (Hotaling et al. 2021). For species in which microsatellite markers have already been developed, and when high-throughput genotyping of microsatellites is an option, choosing between SNPs or microsatellite genotypes remains a relevant decision. High-throughput genotyping of microsatellite amplicons may provide a transitional solution (De Barba et al. 2017; Pimentel et al. 2018; Roy et al. 2021). Microsatellite genotyping takes the PCR products from potentially hundreds of microsatellite primers across hundreds of individuals and sequences repeats that can be bioinformatically sorted with barcodes. In addition to the ability to use previously developed markers that can be compared to older studies and genotype many individuals at a time, this approach directly sequences microsatellite repeats and avoids the meticulous nature of hand scoring peaks from capillary electrophoresis (Guichoux et al. 2011). This approach was pioneered in human forensics for bulk genotyping of short tandem repeats (Fordyce et al. 2011) and subsequently expanded into non-model organisms (Vartia et al. 2016; Darby et al. 2016; De Barba et al. 2017; Pimentel et al. 2018; Salado et al. 2021). The development of bioinformatics tools to process these data has shortly followed their expanded use in ecology and conservation (Barbian et al. 2018; Lepais et al. 2020; Roy et al. 2021).

In this study, we use previously designed microsatellite primers for high-throughput genotyping of a threatened frog species, the Chiricahua Leopard Frog (*Lithobates chiricahuensis*). Like many amphibians of conservation concern, a combination of habitat fragmentation, predation and competition from invasive species, and the fungal pathogen *Batrachochytrium dendrobatidis* (Bd) has resulted in a drastic reduction in the number of Chiricahua Leopard Frog populations across their southwestern US range (USFWS 2007; Rorabaugh and Sredl 2014). Management of the species through releases and translocations began at a small scale in the 1990s, expanded after the species was federally listed as threatened in 2002, and again when the Recovery Plan was published in 2007 (USFWS 2007; Rorabaugh et al. 2008; Sredl et al. 2011). To date, more than 40,000 frogs and tadpoles and hundreds of egg masses have been released or translocated to wild sites in the species’ historical range in Arizona and New Mexico (Hossack et al. 2022) with the intent of starting new populations, augmenting existing populations, and ultimately providing Chiricahua Leopard Frogs with the opportunity to evolve Bd-resistance in the wild.

To meet the recovery goal of delisting, the frog must reach a population level and have sufficient habitat distributed throughout its historical range to provide for the long-term persistence of metapopulations in each of eight recovery units (RUs), even in the face of local extirpation. Conserving the frog within each RU ensures that when recovered, it will be well-distributed and threats will be lessened or alleviated throughout its historical range. If it is conserved and well-distributed within its historical range, then it will no longer be threatened. This strategy and the implementation of recovery actions address the needs of each RU and focus on management areas (MAs), which are areas within RUs with the greatest potential for successful recovery actions and threat alleviation. MAs contain extant populations or sites where habitats will be restored or created, and populations of frogs established or re-established. While genetic diversity of Chiricahua Leopard Frogs is not a recovery criterium, the Recovery Plan outlines the need for a genetic management plan to maintain or enhance genetic diversity within each of eight

Recovery Units (RU; USFWS 2007). With no genetic management plan in place, genetic management has broadly focused on maintenance of genetic preservation or representation within and among the RUs, watersheds, management areas, or specific mountain ranges (McCall et al 2018). This approach has largely limited genetic mixing across these bounds to preserve perceived genetic integrity and local adaptation of populations. Because of the significant rangewide declines, a conservative approach towards genetic management, and captive propagation inherently involving small numbers of breeders representing a RU, many populations have experienced bottlenecks or founder effects.

Several genetic studies have included Chiricahua Leopard Frogs Frogs (Goldberg et al. 2004; Hillis and Wilcox 2005; Hekkala et al. 2011), but the focus of these projects has primarily been to delineate lineages of the species from other members of the phenotypically-similar clade of North American leopard frogs. Chiricahua Leopard Frogs remain in need of genetic assessment, especially given the documented bottlenecks associated with reintroductions and captive breeding. In this study, we used a large set of microsatellite loci to compare populations with different management histories, cluster populations across the species’ range into related groups, and measure the change in allelic richness of a single, heavily managed and bottlenecked population over the course of a decade. Finally, we use simple permutations of our total number of loci to estimate the influence of the number of microsatellite loci used on our estimates.

## Methods

### Sampling and tissue collection

For the rangewide genetic assessment, we selected 62 samples from individuals across the range in an effort to maximize the diversity of populations and management strategies used at individual sites. For most individuals, a toe-clip or tail-clip was collected by state agency personnel, and tadpoles were randomly sampled across the habitat to reduce the chances of collecting siblings. Chiricahua Leopard Frog genetic lineages are represented across a spectrum of “populations”, with some genetic stock represented in functioning metapopulations in the wild, and other genetic stocks only represented in frogs held in captive or semi-captive habitats. We wanted representation from as much of the remaining genetic stock as possible, so we broadly defined “populations” to include metapopulations (a grouping of occupied, wild sites connected via dispersal); isolated, occupied wild sites; and genetic lineages represented in captive colonies. To compare diversity metrics across a spectrum of management histories, we defined nine management categories, ranging from populations with no or little history of translocations, to populations heavily managed and only found in captive or semi-captive colonies (Table 1).

**Table 1.** Management histories of Chiricahua Leopard Frog (*Lithobates chiricahuensis*) populations used for rangewide genetic diversity assessment. Categories are listed in order of least managed to most managed.

| Management Category | Definition |
| --- | --- |
| No Management | No history of management through translocations or isolated site with no known population origin. Emigration and immigration occur or presumed to occur across a functioning metapopulation |
| Unmanaged/Declining | Population has had no translocations, but is small, isolated, or declining |
| Minimally Managed | Multiple wild to wild translocations have occurred and emigration and immigration occur or are likely to occur within metapopulation |
| Captive Propagation | Metapopulation is augmented with offspring of breeders from captive facility; emigration and immigration occur within management area |
| Heavily Managed | Wild to wild translocations and/or captive propagation releases have occurred within population; natural metapopulation functioning unlikely |
| Refugia Site | Genetic stock represented in an isolated population following wild population extirpation |
| Few Founders | Population initially started with few, closely related individuals |

For the long-term cohort sampling, we selected 34 samples from the Southwestern Research Station site (SWRS; Cave Creek, Northern Chiricahua Mountains MA, RU 3) to take advantage of longitudinal tissue samples collected since 2011. Chiricahua Leopard Frogs were extirpated from the Chiricahua Mountains by the 1990s (USFWS 2007), but a reintroduction effort began in the area in 2011 with the creation and restoration of lentic habitats along Cave Creek, a lotic system from which the holotype of the species was described in 1979 (Platz and Mecham, 1979). At the SWRS site, 13 PIT-tagged frogs were released in 2011, originally sourced from another RU 3 population in an adjacent MA (Swisshelm Mountains) at Leslie Creek at Leslie Canyon National Wildlife Refuge (LCNWR); four of these 13 frogs were recaptured in 2013 to get tissues. Another six adult frogs were removed from the ranarium head starting facility in 2011 by Arizona Game and Fish Department (AZGFD) biologists after a die-off event (N = 10 total from the 2011). We expect this 2011 cohort of samples to include “founder” animals or potentially the first generation of SWRS founders. We collected tissue samples from 65 and 24 Chiricahua Leopard Frogs at SWRS in 2013 and 2018, respectively. From the 2013 and 2018 collections, we selected a balance of tissues between adult (male and female) and juvenile animals in the case that genetic assignment could take place across only a few generations. In 2013 and 2014, additional introductions of frogs from the same source (LCNWR) occurred at four other small ponds that sit along a 6 km stretch of Cave Creek, which is within known dispersal distance for the species within a monsoon season (Hinderer et al. 2017). Because of the proximity of these occupied habitats, they likely represent a metapopulation, where frogs from the other ponds likely immigrated periodically into the SWRS site using Cave Creek as a dispersal route. Additionally, an emergency frog salvage was performed in 2017 when Leslie Creek temporarily dried, and 52 larvae, 6 juvenile, and 42 adult frogs were released from Leslie Creek to the SWRS site.

### Screening Microsatellite Loci

We obtained the primer sequences and associated metadata for microsatellite loci that were identified from Chiricahua Leopard Frog genome sequencing by the Cornell University Evolutionary Genetics Core Facility. Microsatellite repeat regions were identified using msatcommander (Rozen and Skaletsky 2000; Faircloth 2008) on sequence data generated from a Illumina MiSeq run (2 x 250bp paired reads) after hybridization to magnetic capture repeat probes (two unique dimers, five unique trimers, seven unique tetramers, and two unique pentamers). The output of msatcommander included more than 11,000 non-duplicated dimeric, trimeric, and tetrameric repeat regions and associated primer pairs. A second run of msatcommander was conducted to focus on sequence lengths 195-205bp and tetramer repeats only (N = 843). Using this list of 843 loci, we removed loci that had duplicate primer sequences, were at the extremes of the coverage distribution, or had outlier repeat lengths. Of 534 microsatellite loci remaining, we divided them into 4-5 groups of ∼25 loci per sequencing reaction, grouping based on the lowest risk of binding to one another (primer dimers) or amplifying inconsistently across all individual DNA samples.

We used the program Autodimer v1.1 (Vallone and Butler 2004) with default settings to identify primer dimers, but the entire set of 534 loci caused resulting groupings to be too large for proceeding to clustering steps so we ran Autodimer twice, once on the first 267 primer pairs (subset1) and once for the second 267 primer pairs (subset2). The output of Autodimer was imported into R (R Core Team 2017), and the lists of potential primer dimers were organized and made into a matrix using the R package tidyr (Wickham et al. 2019). Then, we used the igraph package (Csardi et al. 2006) to optimally cluster primer pairs based on the identification of primer dimers from Autodimer. The “louvain” option produced three reasonably-sized groups of primers from the first subset of loci (A = 111, B = 118, C = 38). These group assignments were exported from R and groups A and B were run through Autodimer again. This process removed an additional 10 primers from group A and 12 from group B. We ordered five sets of primers (29-35 primer pairs/set) from each group (GroupA_order1 = 29 primer pairs, GroupA_order2 = 30, GroupA_order3 = 30, GroupB_order1 = 35, GroupB_order2 = 35). Nexterra tags would need to be added to these primers before high-throughput sequencing, so these tags were added before ordering in order to best predict which primer pairs would amplify well in combination when in the sequencing lane. The Nexterra index tags were added to the 5’ end of each forward and reverse primer (F-primer tag = TCGTCGGCAGCGTCAGATGTGTATAAGAGACAG, R-primer tag = GTCTCGTGGGCTCGGAGATGTGTATAAGAGACAG).

Each pair of primers (159 total) was amplified individually using three different Chiricahua Leopard Frog DNA samples (Table 2). We extracted DNA from toe tissues preserved in ethanol using a Qiagen DNeasy Blood and Tissue Kit. To increase DNA concentration, we decreased the final elution volume by 50% of the recommended volume. Each combination of DNA sample and primer pair was amplified using 5ul BioMix Red (Bioline), 0.5ul forward primer, 0.5ul reverse primer, 4.5ul of water, and 0.5ul of DNA. PCR conditions were 94°C for 1 minute, 35 amplification cycles (94°C for 45s, 56°C for 45s, 72°C for 30s), and an annealing step of 72°C for 10 minutes. We visualized PCR products using ethidium bromide staining on 2% agarose TBE gel that ran at 120V for 20-30 minutes. If a locus failed to amplify in more than a single individual, it was removed from further consideration, and the intensity and clarity of each band was recorded. After this initial screening, we made 2uM multiplex mixes for the best 25 primer pairs in each of the five orders (5 multiplex mixes containing 25 primer pairs each). We used these mixes to perform a multiplex PCR using the Qiagen Multiplex Kit following the recommendations for PCR conditions when using many microsatellite loci. These PCR products were visualized on a gel to confirm the expected size of the reactions (∼200bp).

**Table 2.** Individual Chiricahua Leopard Frog (*Lithobates chiricahuensis*) samples used to screen 159 microsatellite loci.

| Lab# | Extraction Date | DNA Concentration (ng/ul) | Recovery Unit | Population | Locality | Date Collected |
| --- | --- | --- | --- | --- | --- | --- |
| LC064 | 10/31/2019 | 10.7 | 3 | Northern Chiricahuas | Southwestern Research Station | 8/3/2013 |
| LC092 | 11/14/2019 | 7.47 | 1 | Sierrita Mountains | Pond 3 | 7/5/2017 |
| LC122 | 2/11/2019 | 11.8 | 8 | Mimbres Chino | Grant County, NM: Martin Canyon (Ash Spring) | --- |

### Sequencing and quality control

We sent multiplex mixes to Cornell University’s Evolutionary Genetics Core Facility to be sequenced using an Illumina MiSeq using 250bp paired reads and ∼15 million reads. The five multiplex mixes described above were amplified with 96 Chiricahua Leopard Frog DNA samples that were extracted from preserved tissues using the same Qiagen DNeasy procedure (Supplementary Table 1).

Sequencing reads were genotyped using the *amplicon.py* script (available here: https://bitbucket.org/cornell_bioinformatics/amplicon/src/master/). The minimum alleles frequency was set to 0.005, the minimum haplotype length was set to 75, and the maximum read count ratio between the two alleles in each sample was set to 30. After running this script, loci were removed where 20% of the data shows missing or low reads and samples were removed that were missing 20% or more of the remaining loci. This resulted in a final set of 91 microsatellite loci that met these requirements and the removal of three individuals (LC132, LC133, and LC156).

### Genetic analyses

We used PGDSpider v2.1.1.5 (Lischer and Excoffier 2012) to convert the haplotype tables produced by the *amplicon.py* script to genepop format and used the R package *diveRsity* (Keenan et al. 2013) to calculate summary statistics. We then used GenAlEx v6.5 (Peakall and Smouse 2012) to confirm these statistics and identify private alleles for each metapopulation. To investigate how many microsatellite loci are necessary to generate accurate statistics of genetic diversity, we repeated the calculations above with R (R Core Team 2017) using random selections of smaller sets of loci (10, 20, 30, 40, 50, 60, 70, 80, 91). Each sampling was conducted 10 times for populations that had equal sample sizes (Apache Tularosa, Three Forks, Lower Gila, Seco, Sierritas).

We performed genetic clustering using a Discriminant Analysis of Principal Components (DAPC) from the R package *adegenet* (Jombart 2008; Jombart et al. 2010). This method uses sequential K-means clustering and reports Bayesian Information Criterion (BIC) to describe clusters of genetically similar individuals without using any prior information. The number of principal components included in the DAPC was determined by first inspecting the results of a cross validation (*xvaldapc* command). For the SWRS cohort data specifically, a genetic principal component analysis (PCA) was performed in order to plot samples in genetic variation space without clustering groups. To test for heterozygosity excess (recent bottleneck) or deficiency (inbreeding), we used the program Bottleneck v.1.2.02 (Cornuet et al. 1999; Luikart and Cornuet 2008) with a T.P.M. mutation model (95% SMM and a variance of 12), 1,000 permutations, and both Sign Tests and Standardized Differences Tests.

## Results

### Population Genetic Structure

The best supported number of genetic clusters based on lowest BIC score using the *find.clusters* command was three, but a solution of two had a similar BIC score, indicating that both solutions are useful descriptions of genetic structure in these frogs. We used 10 principal components for input into the DAPC and retained both discriminant functions. We found populations broadly split into East and West groups. The three cluster solution separated the populations from southeastern Arizona and groups central Arizona populations with those from southern Arizona and Mexico (Figure 1, 2). The only population with <0.99 probabilities of assignment was frogs representing a genetic mix of the Gentry and Buckskin MAs, which was only assigned to cluster two with a 0.62 posterior probability.

**Figure 1.**
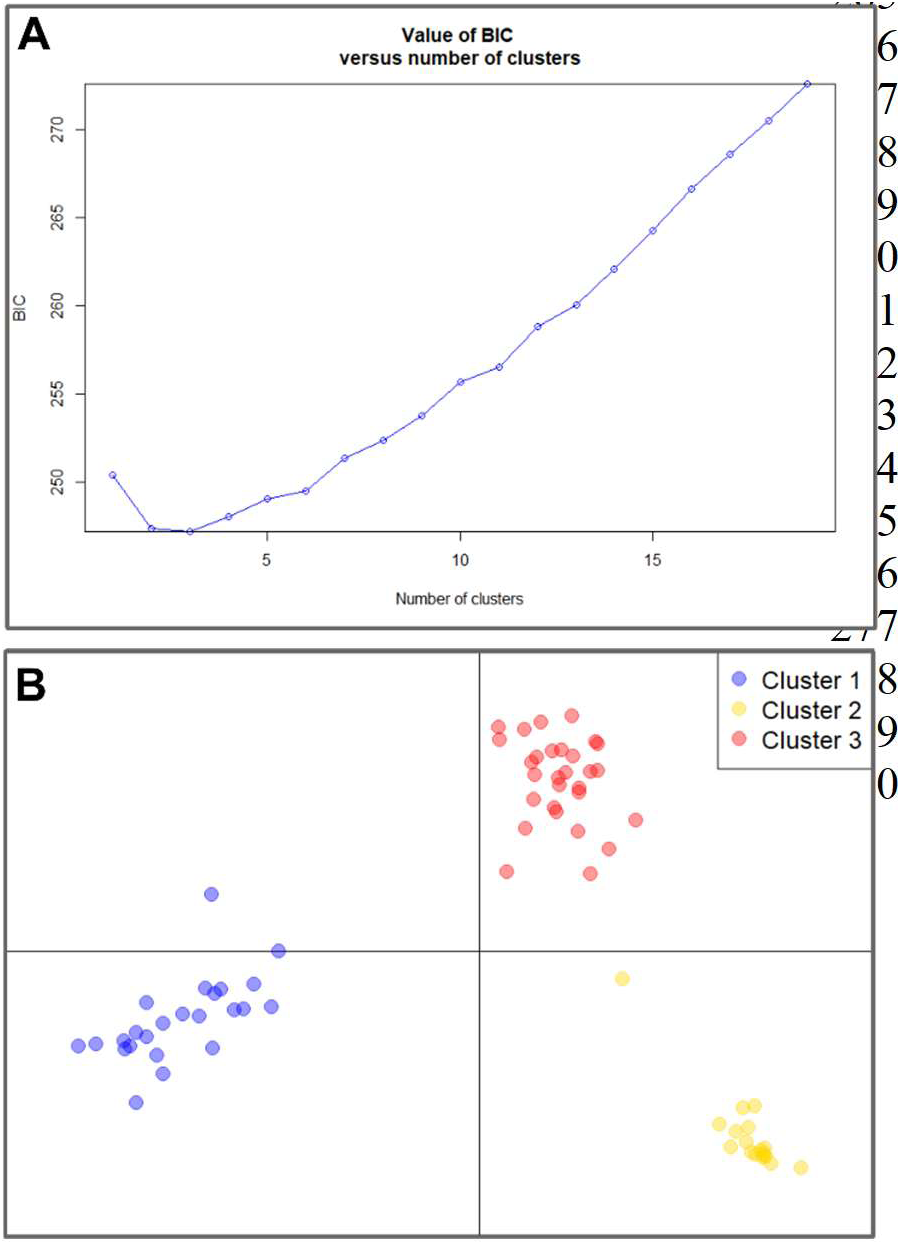
Results of a Discriminant Analysis of Principal Components (DAPC) to find optimal genetic clustering of populations. Panel A shows the results of the *find.clusters* procedure in *adegenet* that compares the statistical fit of different numbers of genetic clusters. The number of clusters with the lowest BIC score is interpreted as the most appropriate clustering solution. Panel B shows how the three clusters are related to one another in variable space. Points represent individual frogs and are colored by their assigned cluster. Points that are closer to one another in space are more similar.

**Figure 2.**
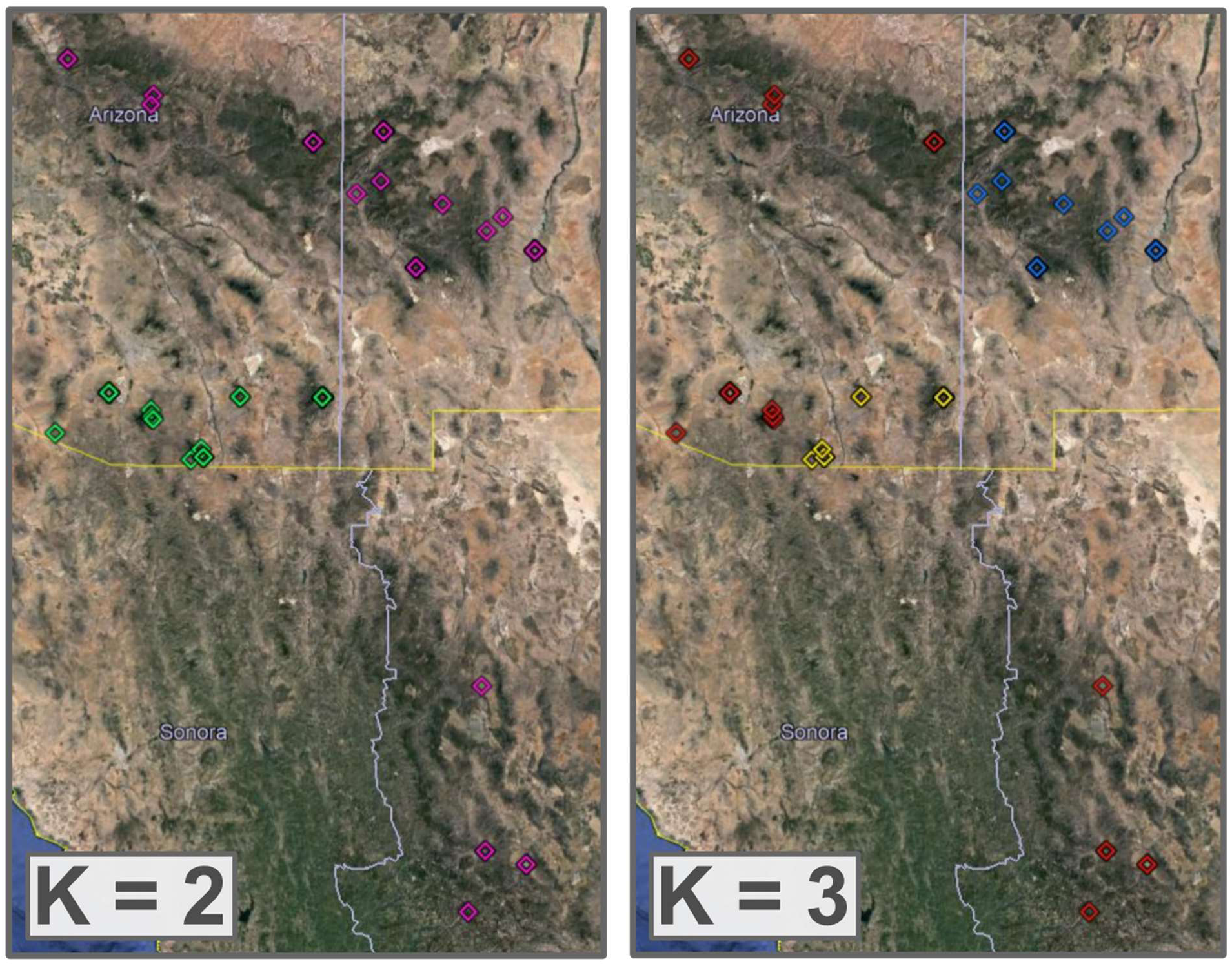
A visualization of the optimal clustering solution with the lowest BIC score (K = 3) and the second lowest BIC score (K = 2). Groups are colored with the same group assignments as in Figure 2.

### Allelic Richness and Inbreeding

A total of 91 microsatellite regions were genotyped after processing, and we used these to assess levels of genetic diversity at both the species’ and population levels. Across all individuals, there were an average of 30.1 alleles/locus (range, 5-64). The mean allelic richness across populations was 3.35 (range = 1.84-5.56; Table 1, Figure 3). All populations were in Hardy-Weinberg equilibrium (*p* > 0.48) except for the Northern Chiricahuas population (*p* < 0.001). The mean number of private alleles (alleles that are only found in one population) was 32.4 (range 8.2-100). Inbreeding coefficients (F_IS_) were only calculated for populations with at least two individuals sampled, and F_IS_ averaged -0.39 (range = -0.65 to -0.16; Table 1).

**Figure 3.**
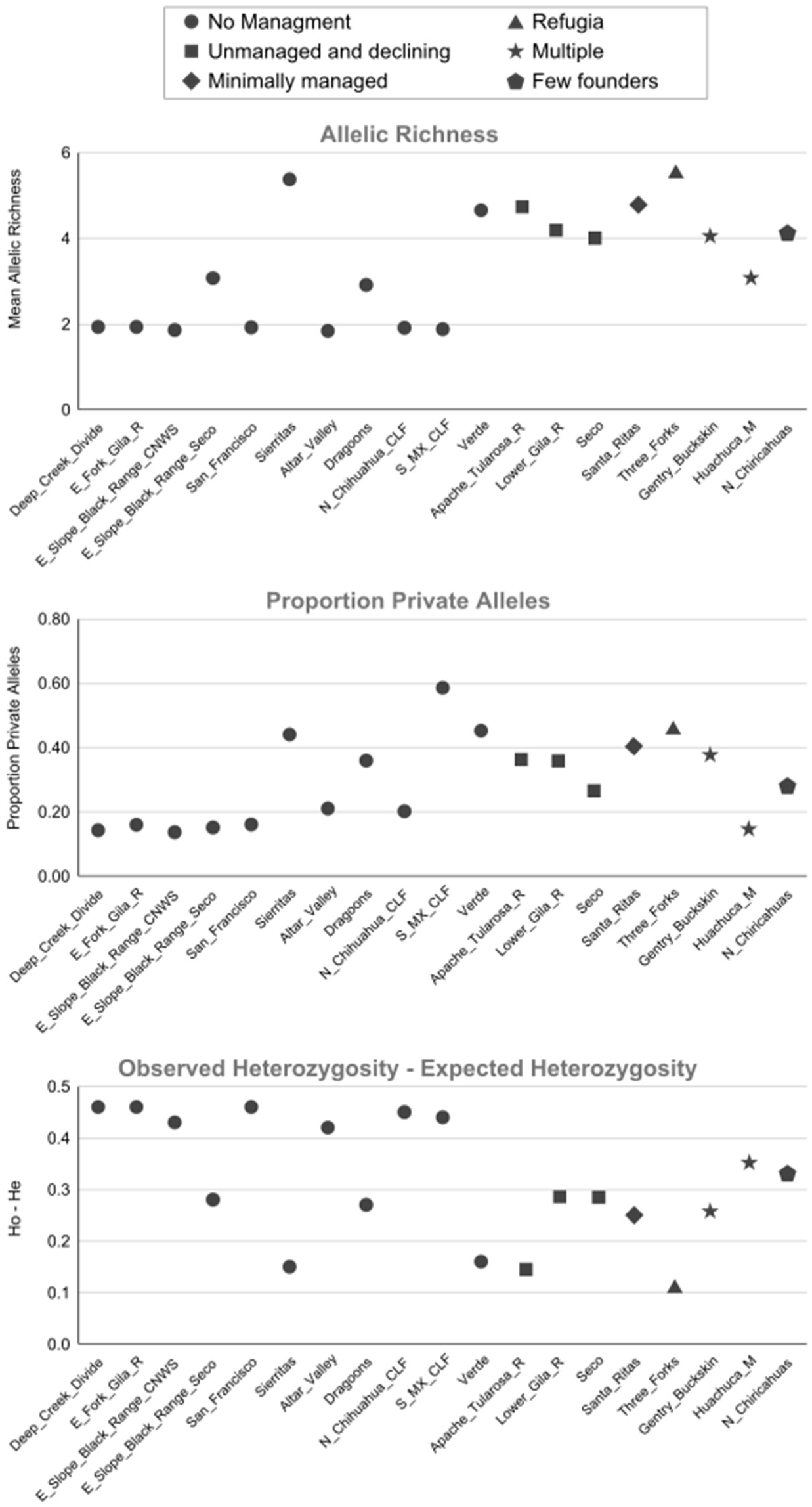
Comparison of allelic richness (top), the proportion of the total alleles that are unique to a population (private alleles; middle), and the difference between observed and expected heterozygosity (bottom) across all populations according to their management histories at the time of tissue collection.

Two populations (Huachuca and Northern Chiricahua Mountains MAs) had significant levels of heterozygosity excess, indicating recent genetic bottlenecks (Table 2). Four populations (Apache Tularosa, Santa Ritas, Sierritas, and Three Forks) had significant levels of heterozygosity deficiency, potentially indicating more long-term inbreeding. The remaining four populations (Gentry-Buckskin mix, Lower Gila River, Seco, and Verde) met the null assumption of heterozygosity-homozygosity equilibrium.

**Table 1.** Genetic summary statistics for 19 Chiricahua Leopard Frog populations and five breeding cohorts of frogs at the Southwestern Research Station (SWRS).

| Grouping | #<br>Samples | # of<br>Alleles | Allelic<br>Richness<br>(average #<br>alleles/locus) | Observed<br>Heterozygosity<br>(H<sub>O</sub>) | Expected<br>Heterozygosity<br>(H<sub>E</sub>) | H<sub>O</sub>-H<sub>E</sub> | p-value<br>(Hardy<br>Weinburg<br>Equilibrium) | Mean<br>F<sub>is</sub> | Total<br>Private<br>Alleles | Proportion<br>Private<br>Alleles |
| --- | --- | --- | --- | --- | --- | --- | --- | --- | --- | --- |
| **Populations** |  |  |  |  |  |  |  |  |  |  |
| Altar_Valley | 1 | 167 | 1.84 | 0.85 | 0.43 | 0.42 | 0.48 | - | 35 | 0.21 |
| Apache_Tularosa_R | 6 | 425 | 4.67 | 0.79 | 0.65 | 0.14 | 1.00 | -0.22 | 151 | 0.36 |
| Deep_Creek_Divide | 1 | 176 | 1.93 | 0.93 | 0.47 | 0.46 | 0.48 | - | 25 | 0.14 |
| Dragoons | 2 | 265 | 2.91 | 0.87 | 0.60 | 0.27 | 1.00 | -0.45 | 95 | 0.36 |
| E_Fork_Gila_R | 1 | 176 | 1.93 | 0.93 | 0.47 | 0.46 | 0.48 | - | 28 | 0.16 |
| E_Slope_Black_Range_CNWS | 1 | 169 | 1.86 | 0.86 | 0.43 | 0.43 | 0.48 | - | 23 | 0.14 |
| E_Slope_Black_Range_Seco | 2 | 279 | 3.07 | 0.89 | 0.61 | 0.28 | 0.99 | -0.46 | 42 | 0.15 |
| Gentry_Buckskin | 4 | 371 | 4.08 | 0.91 | 0.65 | 0.26 | 1.00 | -0.41 | 139 | 0.37 |
| Huachuca_M | 5 | 279 | 3.07 | 0.88 | 0.53 | 0.35 | 0.16 | -0.65 | 41 | 0.15 |
| Lower_Gila_R | 6 | 377 | 4.14 | 0.90 | 0.62 | 0.28 | 0.99 | -0.45 | 131 | 0.35 |
| N_Chihuahua_CLF | 1 | 174 | 1.91 | 0.91 | 0.46 | 0.45 | 0.48 | - | 35 | 0.20 |
| N_Chiricahuas | 12 | 374 | 4.11 | 0.89 | 0.56 | 0.33 | 0.00 | -0.59 | 104 | 0.28 |
| S_MX_CLF | 1 | 171 | 1.88 | 0.88 | 0.44 | 0.44 | 0.48 | - | 100 | 0.58 |
| San_Francisco | 1 | 175 | 1.92 | 0.92 | 0.46 | 0.46 | 0.48 | - | 28 | 0.16 |
| Santa_Ritas | 5 | 436 | 4.79 | 0.93 | 0.68 | 0.25 | 1.00 | -0.38 | 184 | 0.42 |
| Seco | 6 | 360 | 3.96 | 0.89 | 0.61 | 0.28 | 0.82 | -0.47 | 94 | 0.26 |
| Sierritas | 6 | 489 | 5.37 | 0.84 | 0.69 | 0.15 | 1.00 | -0.21 | 215 | 0.44 |
| Three_Forks | 6 | 506 | 5.56 | 0.84 | 0.73 | 0.11 | 1.00 | -0.16 | 230 | 0.45 |
| Verde | 4 | 423 | 4.65 | 0.86 | 0.70 | 0.16 | 1.00 | -0.23 | 191 | 0.45 |
| **SWRS Cohorts** |  |  |  |  |  |  |  |  |  |  |
| 2011_Adults | 10 | 554 | 6.09 | 0.85 | 0.66 | 0.19 | 1.00 | -0.29 | 280 | 0.51 |
| 2013_Adults | 8 | 412 | 4.53 | 0.87 | 0.61 | 0.26 | 0.62 | -0.42 | 141 | 0.34 |
| 2013_Juveniles | 4 | 294 | 3.23 | 0.88 | 0.58 | 0.30 | 0.85 | -0.53 | 64 | 0.22 |
| 2018_Adults | 9 | 343 | 3.77 | 0.89 | 0.56 | 0.33 | 0.00 | -0.59 | 76 | 0.22 |
| 2018_Juveniles | 3 | 239 | 2.63 | 0.90 | 0.53 | 0.37 | 0.25 | -0.70 | 16 | 0.07 |

**Table 2.** Summary of results from the program Bottleneck (Cornuet et al. 1999; Luikart and Cornuet 2008). Two tests of statistical significance for heterozygosity excess (indicating a recent genetic bottleneck) or heterozygosity deficit (indicating ongoing inbreeding) were used, the Sign Test and the Standardized Differences Test. Rows are bolded to indicate populations that rejected the null hypothesis of heterozygosity-homozygosity equilibrium after 1000 permutations, and are therefore likely suffering a bottleneck.

| **Population** | **#**<br>**Individuals** | **mean #**<br>**of alleles** | **mean**<br>**proportion of**<br>**heterozygotes** | **# loci with**<br>**heterozygosity**<br>**Excess ($H_e$)** | **Expected #**<br>**loci with**<br>**heterozygosity**<br>**Excess ($H_{eq}$)** | **p-value**<br>**(Sign Test)** | **T2**<br>**value** | **p-value**<br>**(Standardized**<br>**Differences**<br>**Test)** | **Bottleneck**<br>**Conclusion** |
| --- | --- | --- | --- | --- | --- | --- | --- | --- | --- |
| Apache_Tularosa_R | 6 | 4.67 | 0.72 | 44 | 53.210 | 0.026 | - | - | Het Deficient |
| Gentry_Buckskin | 4 | 4.08 | 0.74 | 60 | 53.090 | 0.068 | 1.460 | 0.072 | Equilibrium |
| Huachuca_M | 5 | 3.07 | 0.59 | 66 | 47.520 | <0.001 | 3.346 | <0.001 | Het Excess |
| Lower_Gila_R | 6 | 4.14 | 0.67 | 50 | 54.680 | 0.179 | -0.624 | 0.266 | Equilibrium |
| N_Chiricahuas | 12 | 4.11 | 0.59 | 56 | 51.120 | 0.173 | -3.598 | <0.001 | Equilibrium/Het<br>Excess |
| Santa_Ritas | 5 | 4.79 | 0.75 | 35 | 51.300 | <0.001 | -6.068 | <0.001 | Het Deficient |
| Seco | 6 | 3.96 | 0.66 | 54 | 54.530 | 0.495 | -0.643 | 0.260 | Equilibrium |
| Sierritas | 6 | 5.37 | 0.77 | 30 | 55.540 | <0.001 | -7.751 | <0.001 | Het Deficient |
| Three_Forks | 6 | 5.56 | 0.82 | 44 | 54.390 | 0.010 | -2.151 | 0.016 | Het Deficient |
| Verde | 4 | 4.65 | 0.81 | 55 | 52.750 | 0.322 | 0.085 | 0.466 | Equilibrium |

As expected, we found that as the number of loci used increased, the smaller the standard deviation became across the randomly selected datasets (Supplementary Figures 1 and 2). However, the number of loci did not drastically influence the calculated values for either Allelic Richness or Inbreeding Coefficient. The largest increases in confidence, as inferred by reductions in standard deviation, broadly take place when using between 20-30 loci.

### Comparison across management histories

The mean allelic richness found in populations with different management histories varied from 2.69 (captive propagation) to 13.77 (unmanaged/declining; Table 4). The proportion of private alleles was highest in unmanaged sites (55%) and lowest in heavily managed sites (18%). Inbreeding coefficients (F_IS_) ranged from -0.63 ± 0.03 SE (Captive Propagation) to -0.06 ± 0.02 SE (Unmanaged).

**Table 4.**
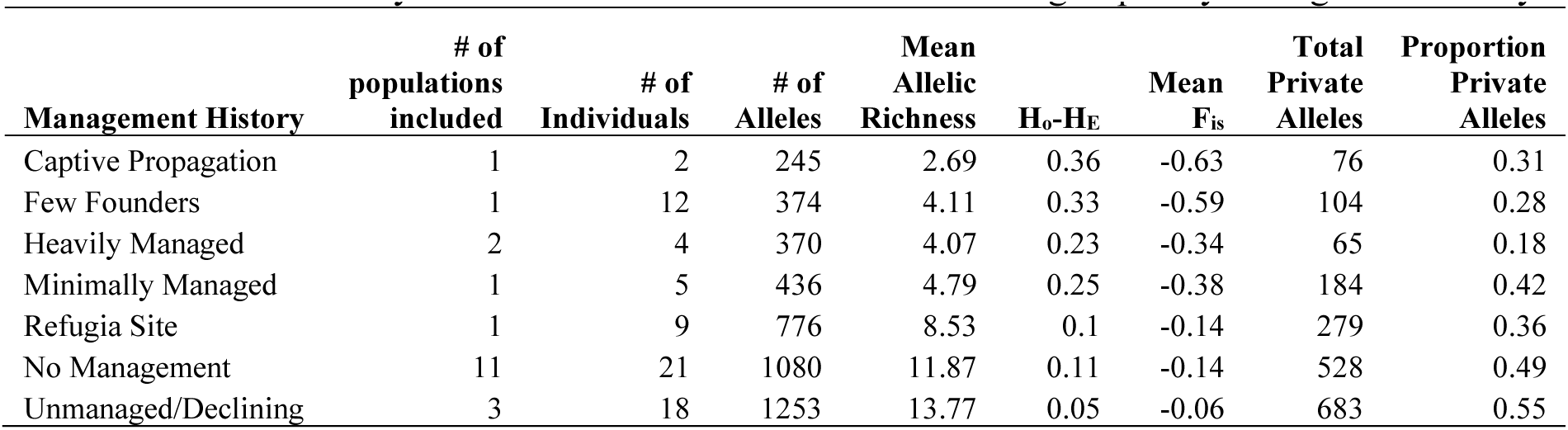
Genetic summary statistics calculated when individuals are grouped by management history.

| Management History | # of<br>populations<br>included | # of<br>Individuals | # of<br>Alleles | Mean<br>Allelic<br>Richness | $H_0-H_E$ | Mean<br>$F_{IS}$ | Total<br>Private<br>Alleles | Proportion<br>Private<br>Alleles |
| --- | --- | --- | --- | --- | --- | --- | --- | --- |
| Captive Propagation | 1 | 2 | 245 | 2.69 | 0.36 | -0.63 | 76 | 0.31 |
| Few Founders | 1 | 12 | 374 | 4.11 | 0.33 | -0.59 | 104 | 0.28 |
| Heavily Managed | 2 | 4 | 370 | 4.07 | 0.23 | -0.34 | 65 | 0.18 |
| Minimally Managed | 1 | 5 | 436 | 4.79 | 0.25 | -0.38 | 184 | 0.42 |
| Refugia Site | 1 | 9 | 776 | 8.53 | 0.1 | -0.14 | 279 | 0.36 |
| No Management | 11 | 21 | 1080 | 11.87 | 0.11 | -0.14 | 528 | 0.49 |
| Unmanaged/Declining | 3 | 18 | 1253 | 13.77 | 0.05 | -0.06 | 683 | 0.55 |

### Comparison across SWRS cohorts

Finally, we compared among SWRS cohorts to provide a temporal view of genetic diversity in the head-started population there. Mean allelic richness linearly decreased from 2011 to 2018 (2011 Adults = 6.09, 2018 Juveniles = 2.63; Table 1). The proportion of private alleles was highest in 2011, indicating that a large proportion of alleles have been lost over time. The 2011, 2013, and 2018 sample years clearly define the clustering of samples into three units via DAPC. Moreover, the 2013 and subsequently 2018 population occupy a more constrained genotypic space relative to 2011, such that 2018 frogs cover only a small fraction of the genotypic space that was present in the 2011 founding (Figure 4B). Over time, the genetic identity of the SWRS population has become more constrained and less diverse.

**Figure 4.**
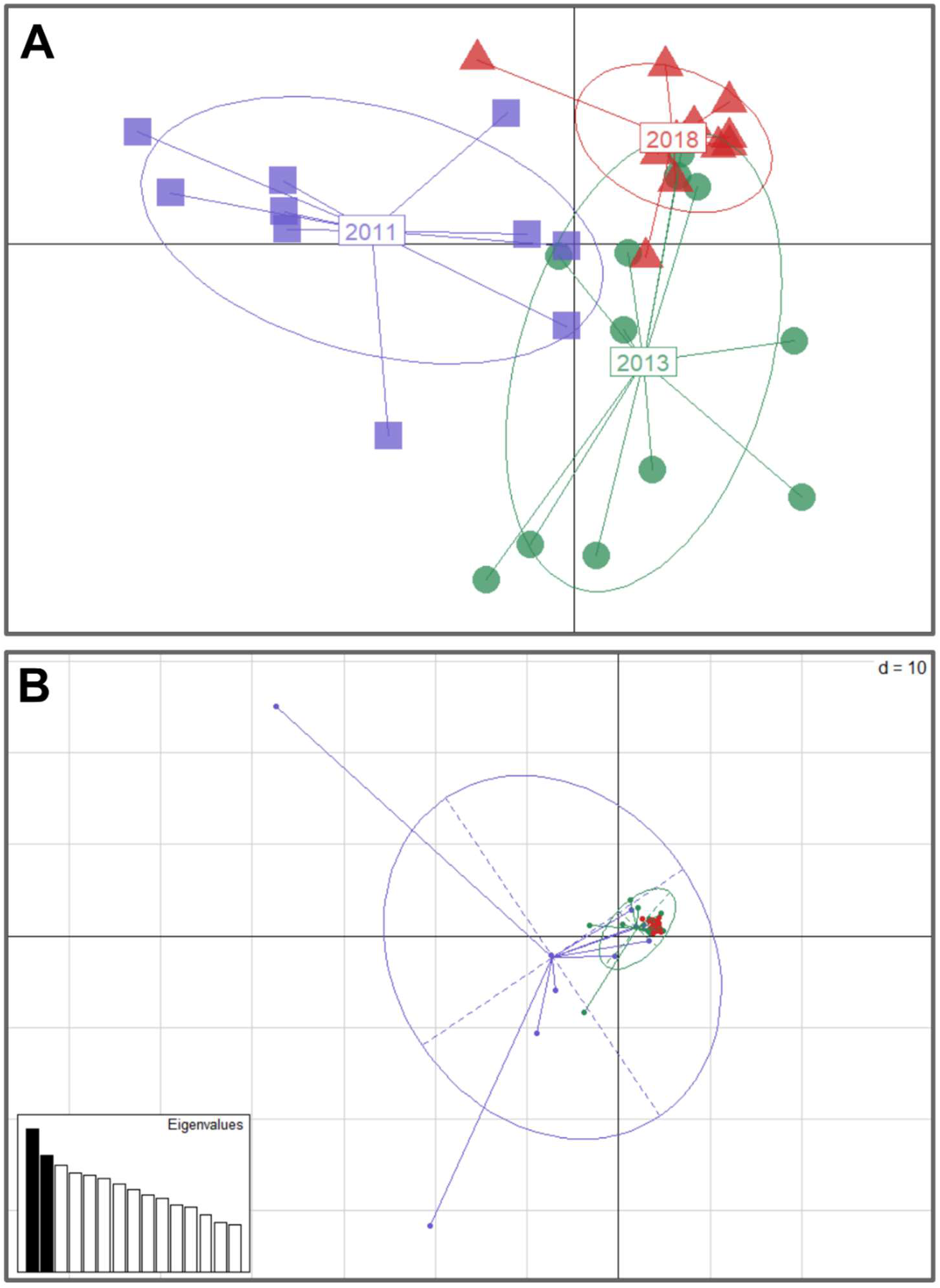
Visualizing the genetic variation from 2011-2018 at the Southwestern Research Station (Cave Creek). Panel A shows the results of a Discriminant Analysis of Principal Components (DAPC) using only the SWRS individuals. Panel B shows the same individuals using a Principal Components Analysis (PCA), which is the first step in the DAPC procedure. This better displays the total variance of the genetic information instead of explicitly attempting to discriminating between different groups. For both panels, the coloring of points is consistent (2011 = purple, 2013 = green, 2018 = red) and different life stages (adults and juveniles) have been collapsed together for clarity. In panel A, the six left-most purple squares coincide with the six 2011 samples.

## Discussion

The Chiricahua Leopard Frog populations represented in this project harbor a much greater amount of genetic variation (>30 unique alleles/locus) than what is portioned into individual populations. Although each population has undergone a different history of management, the results support a relatively recent isolation of populations that may or may not be concurrent with large reductions in population sizes.

While all populations carried some level of genetic uniqueness, as measured by the proportion of private alleles in the total pool of alleles, each population was not different enough to be statistically supported as a unique genetic cluster (DAPC analyses). The best-supported solution is to partition the 19 populations into three broad clusters, one broad (from central Arizona to Mexico) and two more geographically distinct (southeastern Arizona Sky Islands and New Mexico). It is difficult to contextualize these genetic clusters, such as which populations more recently share a common ancestor, without more evolutionary information that is outside the scope of the generated data. Additionally, the sampling gap between populations in New Mexico and Mexico may be strongly influencing the result.

Population genetic estimates may help contextualize the difficulty of identifying genetic clusters. We observed values of observed heterozygosity greater than expected across all populations, and an increase in H_o_-H_e_ over time in the SWRS population. Recent and drastic reductions in population sizes can result in an excess of heterozygous genotypes, and heterozygosity is predicted to decrease more slowly over time than allelic richness (Allendorf 1986, Allendorf et al. 2014). Another mechanism that can produce signals of excess heterozygosity is a scenario where two divergent populations are suddenly mixed with one another, but managers have not purposefully mixed populations from different managements units for fear of outbreeding depression. In populations that have undergone active augmentation by conservation agencies, this could be the case. However, the management strategies for the Chiricahua Leopard Frog are currently focused on maintaining the genetic integrity of historical populations.

It is necessary that each metapopulation included in this project be viewed through the experience of those personnel most informed about the specific history of those sites (year-to-year variation, changes in habitat, introduction of new animals/larvae). Broadly, it is unsurprising that sites classified as unmanaged, natural, or refugia have much greater allelic richness than those that are heavily managed or involved in captive propagation. Managed sites (those with frequent history of moving frogs) show lower genetic diversity than unmanaged populations. Lower genetic diversity could be a result of management or an artifact of populations that required management. In the case of the Chiricahua Leopard Frog, the latter is a more realistic scenario as heavily managed sites are those that conservation personnel have judged as requiring intervention due to combinations of population decline, disease, and habitat alteration. The sites classified as unmanaged, natural, or refugia have relatively low allelic richness when considered individually (Figure 3), but these sites combined have the highest proportion of private alleles. Analyses using DNA sequence data could be paired with the results from this project to determine which populations could be mixed with one another to recover genotypes that are the most historically representative. This would require higher resolution of the genetic variation from sites that may act as either donors or receivers of new breeding individuals.

One example of a well understood and documented set of re-introductions is the population at the SWRS site, which has been carefully documented and regularly sampled since the re-introduction of just 13 animals from Leslie Canyon in 2011 (the majority of which were genotyped here). Although frogs were also released into other ponds along Cave Creek in the following years, likely resulting in dispersal up and down Cave Creek and therefore immigration into the SWRS sites, the source for all the releases was a single, isolated population on Leslie Canyon National Wildlife Refuge. Without any other sources of frogs outside of this single source, the decline in genetic variation over time observed there is the result of a drastic bottleneck, subsequent population growth, and inbreeding. The observed reduction in genetic diversity across our longitudinal samples at SWRS separated by only a few years is bleak, especially as these habitats have been supplemented by constructed breeding habitat and aggressively monitored. Further, this population is predicted to continue to lose genetic variation over time, as the loss of heterozygosity catches up to the initial loss of alleles.

To make the inferences above, we effectively genotyped many microsatellite loci using a high-throughput strategy. This approach is relatively new and can provide significantly more accurate inferences compared to capillary-based sequencing of 10-20 microsatellite markers (De Barba et al. 2017, Pimentel et al. 2018). Compared to other methods for generating many loci for a given study species, such as reduced representation approaches or shotgun genome sequencing, microsatellites still provide more information per locus at the cost of greater upfront design cost and preliminary laboratory work. The permutation analyses in this study provide a unique perspective compared to other empirical comparisons of microsatellite and SNP data. Broadly, other comparisons are most often between a static number of microsatellites and a static number of SNPs from reduced-representation approaches (*reviewed in* Sunde et al. 2020; Zimmerman et al. 2020; Hauser et al. 2021). Prior work to our knowledge has not used many (>30) microsatellite loci to investigate the number necessary to maximize the confidence in estimates of population parameters. Our results align with other work that suggests ∼20 microsatellite loci are broadly comparable to RADseq data (Zimmerman et al. 2020). However, the conclusions are highly dependent on the biological question and system under investigation (Ackiss et al. 2020). In addition to species for which many microsatellite loci have already been designed and screened, this strategy would be most efficient in species with relatively higher amounts of standing genetic variation, as 91 loci were far more than enough to accurately estimate the population genetics statistics in this study. The number of loci genotyped in this project additionally does not make up for a lack of sampling from some populations, and these populations with low sample sizes (two or fewer individuals) had to be excluded from some analyses. However, these individuals are important for understanding the overall patterns of population genetic structure and broad-scale demographic history across the range of Chiricahua Leopard Frogs.

For the broad conservation of the Chiricahua Leopard Frog, this type of genetic surveillance is important to both understand the degree of remaining genetic variation in the species’ range and inform potential future management decisions. To date, managers have worked to keep the genetic lineages of metapopulations preserved and limited mixing to only between nearby populations. Compared to more dire wildlife conservation scenarios in which only a small fraction of the genetic diversity remains in a handful of populations, Chiricahua Leopard Frogs appear to harbor a broad allelic diversity across their management units. Whether of not these conservation genetic scenarios warrant preemptive mixing of populations to maximize genetic diversity instead of continuing to preserve the integrity of small, isolated populations is an ongoing discussion (Powell 2022, Ralls et al. 2018). While there are still few examples of this approach, recent data-driven reintroductions have been successful for other frog species (Knapp et al 2023). Future work should focus on defining a genetic management plan that resolves the distinctness of genetic clusters and can serve as the basis for demonstrating the potential of outbreeding depression (Bell et al. 2019, Hedrick and Fredrickson 2010; Senn et al. 2014). Unlike other conservation scenarios in which only a few populations or even individuals remain, these actions are still viable in Chiricahua Leopard Frogs due to the consistent actions of managers that have helped prevent local extinctions and keep the species within a large part of its historical range.

## Acknowledgements

This work was made possible by funding from the Phoenix Zoo and Arizona Game and Fish Department. We thank Steven Michael Bogdanowicz (Cornell University) for providing additional assistance during marker development and genotyping. We thank Aaron Cajero, Cat Crawford, Abi King, Bill Radke, and Mike Sredl, who salvaged frogs for Leslie Canyon National Wildlife Refuge and provided the source of frogs at SWRS. We thank former and current leaders at the AMNH SWRS, Dawn Wilson and Geoff Bender for supporting frog propagation and habitat creation at SWRS and for supporting the tissue collections on site. We also thank the 2013 and 2018 participants in the AMNH’s “Field Herpetology of the Southwest” course for their participation in the collection of animals and tissues. Broad sampling assistance was provided by Bobby Arnold, Eric Wallace, Bruce Christman, Randy Jennings, Tim Snow, David Hall, Mike Sredl, Abi King, Cody Mosley, Christina Akins, Shaula Hedwall, Cassidi Cobos, and Melanie Culver.

---

**Supplementary Table 1.**
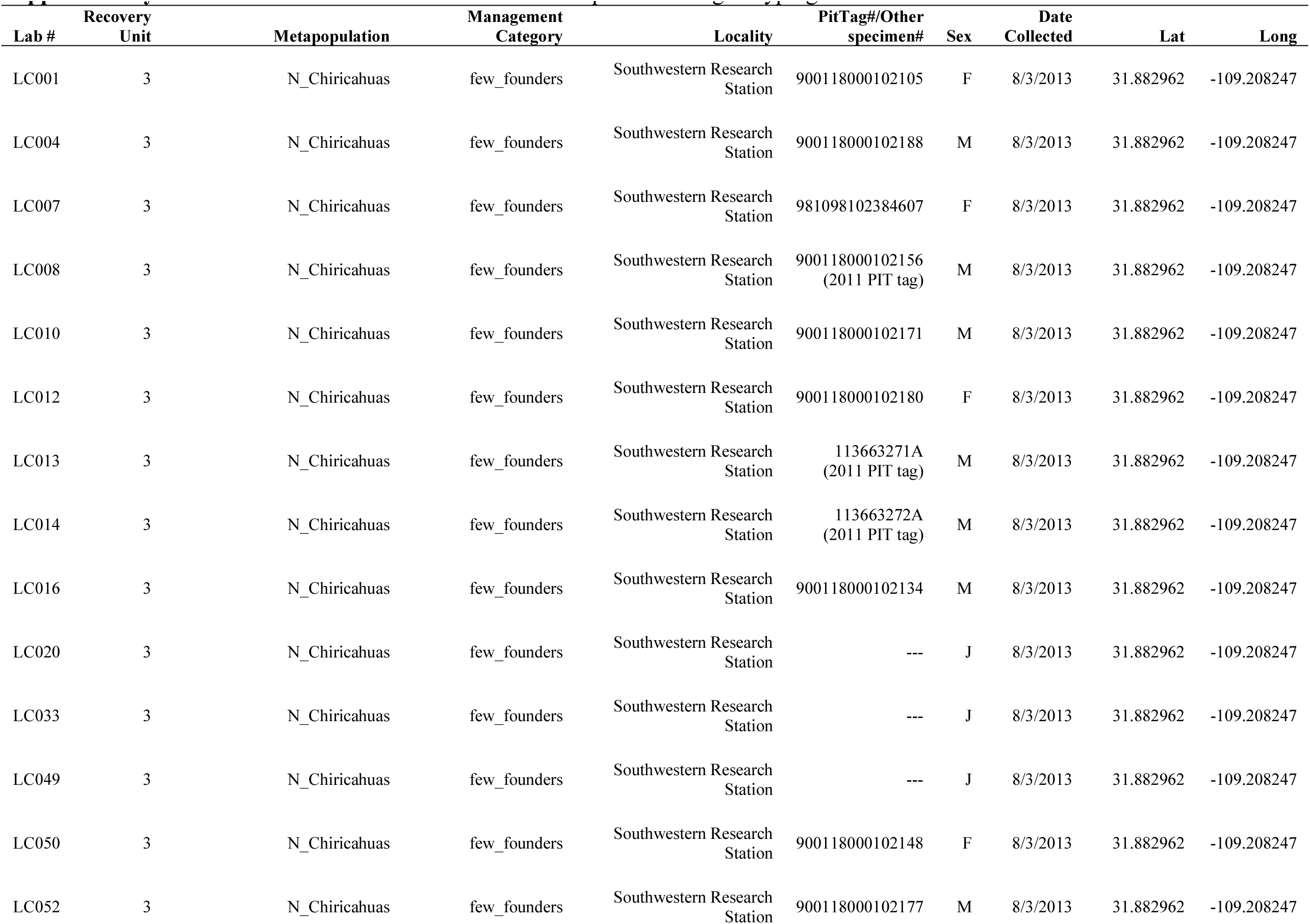
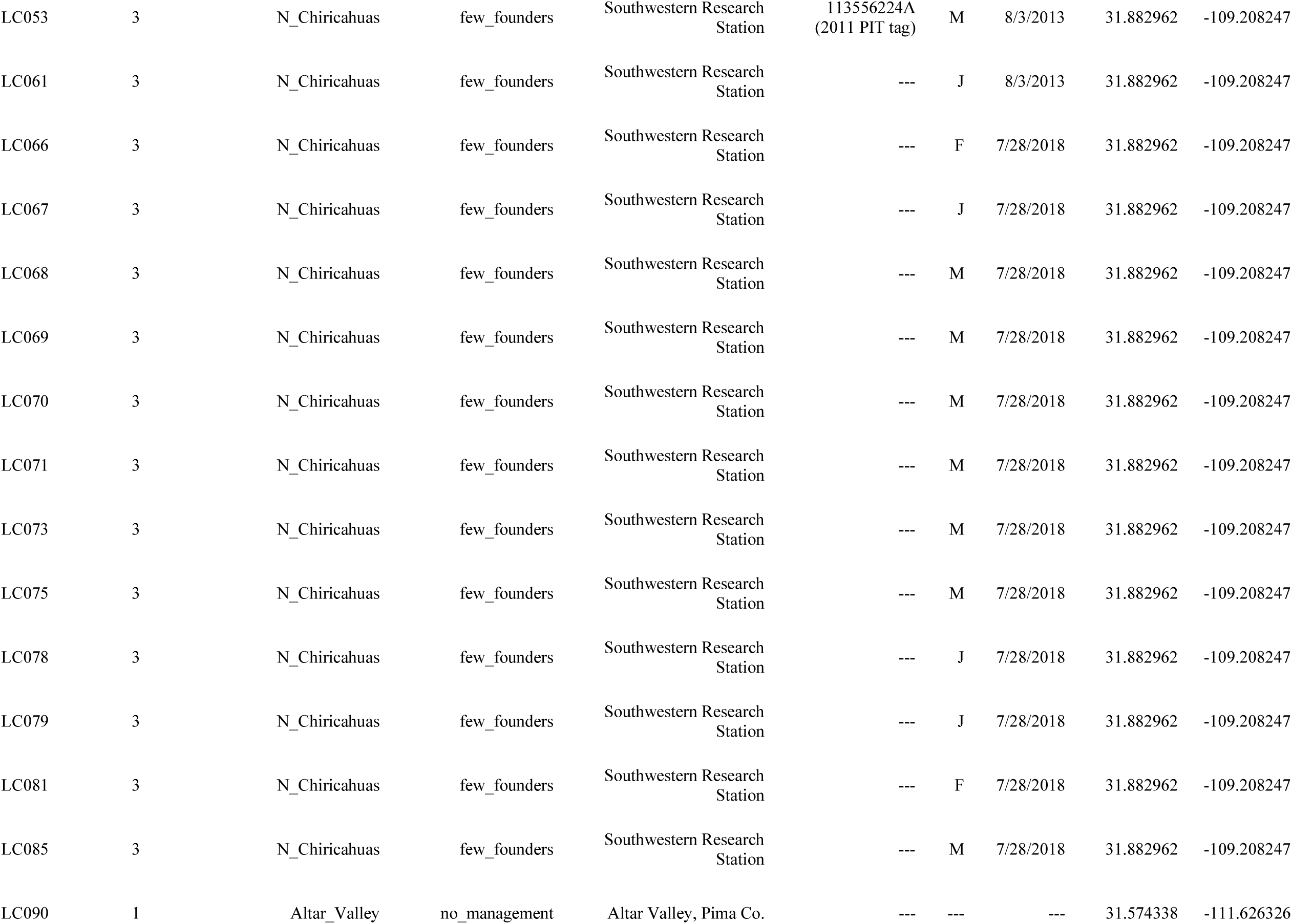
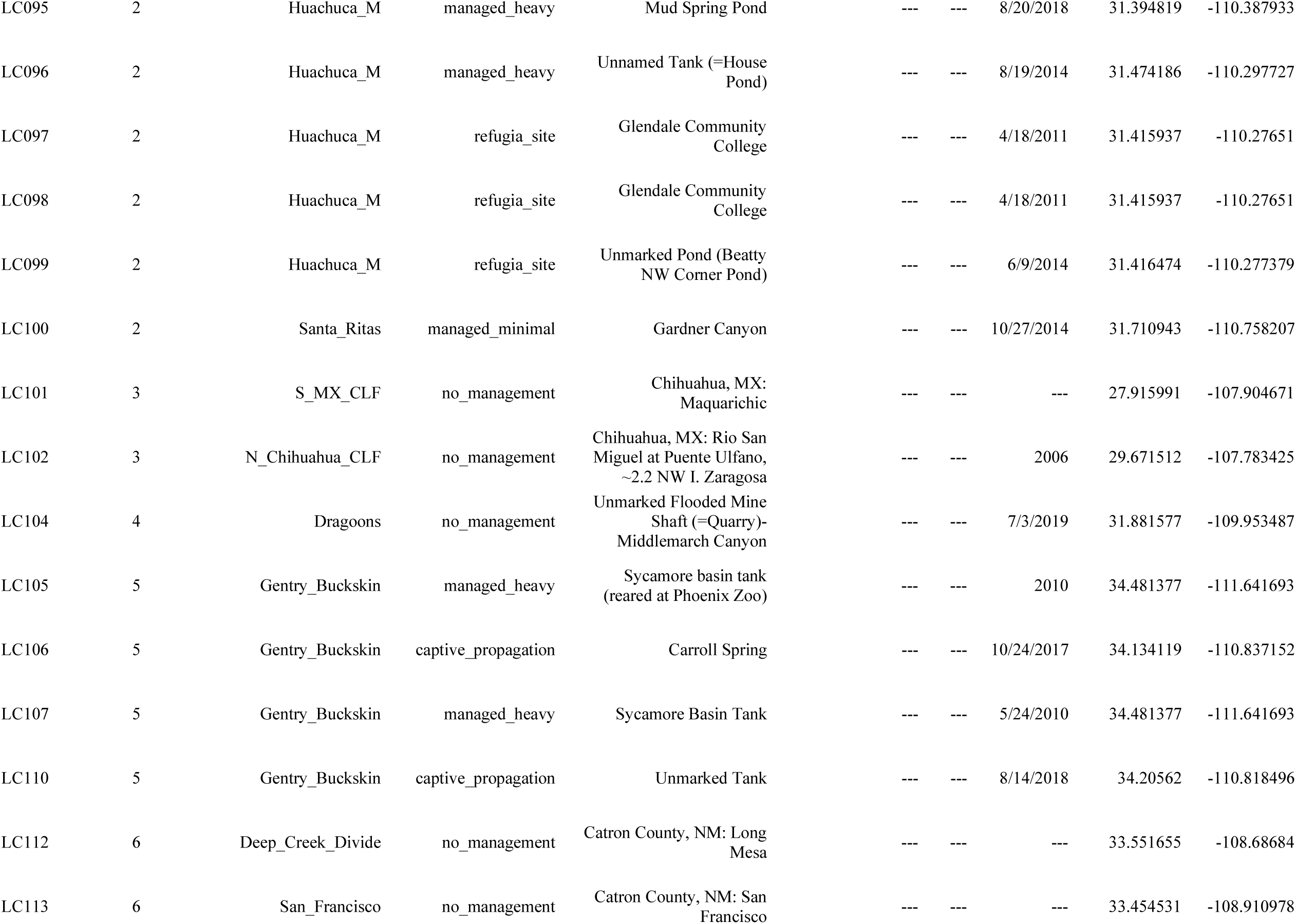
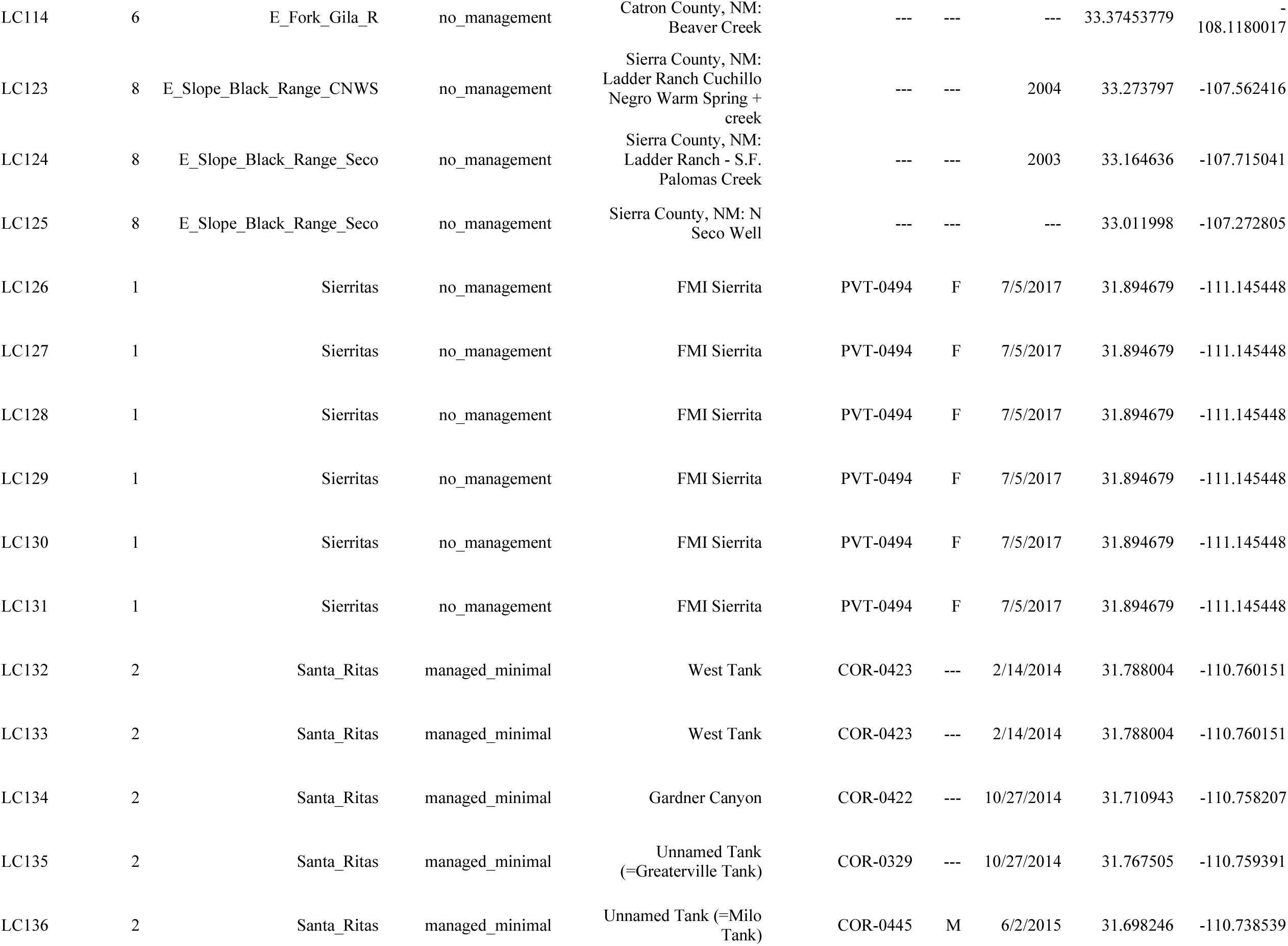
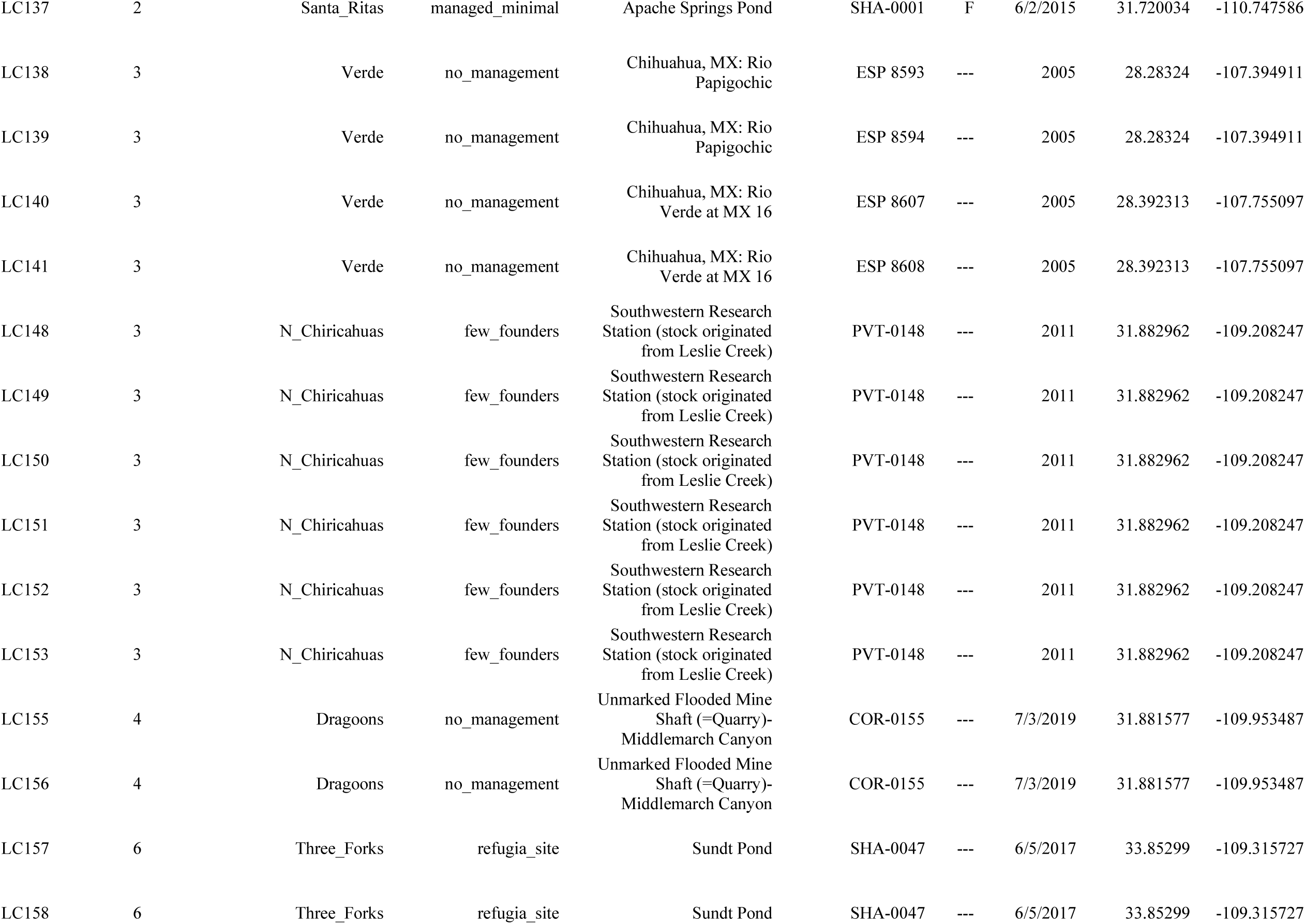
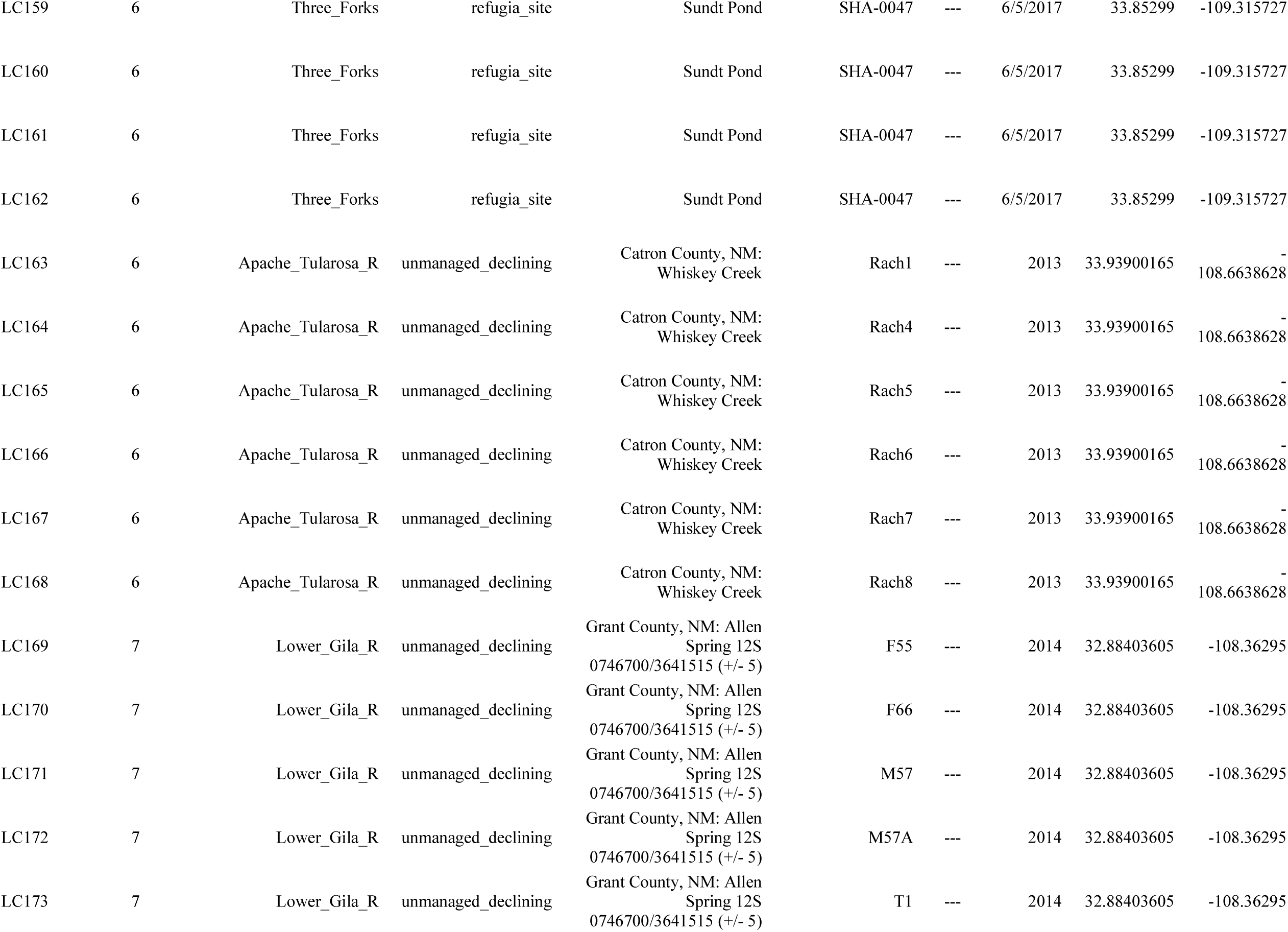
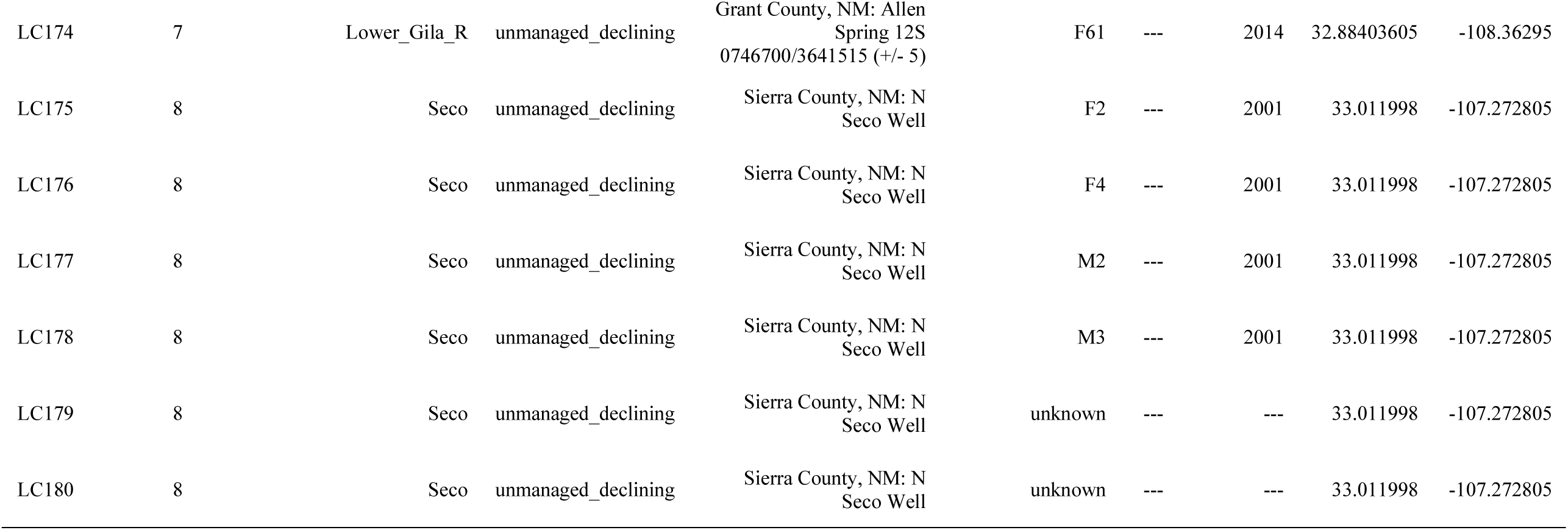
List and associated metadata for samples sent for genotyping.

**Supplementary Table 2.** Summary statistics for all 91 microsatellite loci that remained after quality control.

| **Locus Name** | **# Alleles** | **Simpson Index (1-D)** | **Nei's Genetic Diversity (H<sub>exp</sub>)** | **Evenness** |
| --- | --- | --- | --- | --- |
| Rchi10193 | 35 | 0.9 | 0.91 | 0.57 |
| Rchi10454 | 16 | 0.74 | 0.74 | 0.57 |
| Rchi10490 | 22 | 0.7 | 0.71 | 0.49 |
| Rchi105211 | 18 | 0.82 | 0.83 | 0.67 |
| Rchi105682 | 13 | 0.8 | 0.8 | 0.66 |
| Rchi10626 | 35 | 0.92 | 0.92 | 0.63 |
| Rchi10633 | 13 | 0.74 | 0.75 | 0.68 |
| Rchi10660 | 53 | 0.95 | 0.96 | 0.62 |
| Rchi10906 | 49 | 0.94 | 0.95 | 0.59 |
| Rchi10945 | 38 | 0.93 | 0.94 | 0.69 |
| Rchi10979 | 29 | 0.83 | 0.84 | 0.51 |
| Rchi11060 | 5 | 0.57 | 0.58 | 0.81 |
| Rchi11364 | 52 | 0.93 | 0.94 | 0.54 |
| Rchi11461 | 60 | 0.94 | 0.95 | 0.48 |
| Rchi11697 | 28 | 0.85 | 0.86 | 0.59 |
| Rchi11994 | 12 | 0.56 | 0.56 | 0.52 |
| Rchi12007 | 49 | 0.95 | 0.96 | 0.67 |
| Rchi12234 | 41 | 0.92 | 0.93 | 0.54 |
| Rchi12240 | 37 | 0.85 | 0.85 | 0.41 |
| Rchi12318 | 46 | 0.93 | 0.94 | 0.57 |
| Rchi12466 | 49 | 0.73 | 0.73 | 0.24 |
| Rchi124668 | 23 | 0.89 | 0.9 | 0.7 |
| Rchi12639 | 19 | 0.88 | 0.89 | 0.75 |
| Rchi12714 | 15 | 0.6 | 0.6 | 0.6 |
| Rchi13256 | 43 | 0.91 | 0.92 | 0.47 |
| Rchi13312 | 49 | 0.93 | 0.94 | 0.56 |
| Rchi13427 | 44 | 0.95 | 0.95 | 0.69 |
| Rchi14002 | 18 | 0.78 | 0.78 | 0.6 |
| Rchi14076 | 30 | 0.91 | 0.92 | 0.65 |
| Rchi14085 | 51 | 0.95 | 0.96 | 0.64 |
| Rchi14208 | 18 | 0.89 | 0.89 | 0.75 |
| Rchi14341 | 16 | 0.72 | 0.73 | 0.54 |
| Rchi14362 | 43 | 0.94 | 0.94 | 0.61 |
| Rchi14374 | 15 | 0.6 | 0.6 | 0.51 |
| Rchi14591 | 29 | 0.88 | 0.89 | 0.57 |
| Rchi15189 | 31 | 0.9 | 0.91 | 0.65 |
| Rchi15384 | 12 | 0.63 | 0.63 | 0.68 |
| Rchi15586 | 42 | 0.9 | 0.91 | 0.5 |
| Rchi16001 | 46 | 0.94 | 0.95 | 0.64 |
| Rchi16030 | 12 | 0.7 | 0.71 | 0.67 |
| Rchi16370 | 37 | 0.93 | 0.93 | 0.64 |
| Rchi16650 | 21 | 0.76 | 0.76 | 0.56 |
| Rchi16682 | 43 | 0.95 | 0.95 | 0.65 |
| Rchi16806 | 8 | 0.62 | 0.62 | 0.7 |
| Rchi17068 | 34 | 0.93 | 0.94 | 0.7 |
| Rchi17135 | 39 | 0.92 | 0.92 | 0.59 |
| Rchi17247 | 28 | 0.75 | 0.75 | 0.43 |
| Rchi17339 | 14 | 0.68 | 0.69 | 0.67 |
| Rchi17597 | 30 | 0.77 | 0.77 | 0.41 |
| Rchi18230 | 50 | 0.92 | 0.93 | 0.48 |
| Rchi18664 | 64 | 0.94 | 0.95 | 0.48 |
| Rchi18758 | 16 | 0.71 | 0.72 | 0.54 |
| Rchi18834 | 57 | 0.93 | 0.94 | 0.49 |
| Rchi18837 | 16 | 0.78 | 0.78 | 0.58 |
| Rchi18991 | 18 | 0.64 | 0.64 | 0.53 |
| Rchi19419 | 6 | 0.6 | 0.61 | 0.81 |
| Rchi19501 | 22 | 0.33 | 0.34 | 0.28 |
| Rchi19605 | 24 | 0.81 | 0.82 | 0.55 |
| Rchi19658 | 19 | 0.85 | 0.85 | 0.63 |
| Rchi19676 | 18 | 0.85 | 0.86 | 0.74 |
| Rchi19832 | 38 | 0.94 | 0.94 | 0.67 |
| Rchi19997 | 36 | 0.74 | 0.75 | 0.34 |
| Rchi20229 | 30 | 0.86 | 0.87 | 0.53 |
| Rchi20662 | 25 | 0.8 | 0.81 | 0.53 |
| Rchi20765 | 50 | 0.94 | 0.95 | 0.61 |
| Rchi20833 | 50 | 0.88 | 0.89 | 0.41 |
| Rchi21321 | 15 | 0.76 | 0.77 | 0.56 |
| Rchi21952 | 16 | 0.6 | 0.6 | 0.6 |
| Rchi22483 | 12 | 0.69 | 0.69 | 0.63 |
| Rchi22519 | 10 | 0.17 | 0.17 | 0.34 |
| Rchi23268 | 25 | 0.82 | 0.83 | 0.53 |
| Rchi24447 | 20 | 0.75 | 0.76 | 0.55 |
| Rchi25013 | 19 | 0.82 | 0.83 | 0.66 |
| Rchi25122 | 38 | 0.81 | 0.82 | 0.4 |
| Rchi25188 | 30 | 0.85 | 0.86 | 0.58 |
| Rchi25249 | 22 | 0.8 | 0.81 | 0.58 |
| Rchi25274 | 43 | 0.9 | 0.9 | 0.51 |
| Rchi25518 | 49 | 0.93 | 0.93 | 0.55 |
| Rchi26050 | 47 | 0.95 | 0.96 | 0.68 |
| Rchi26129 | 27 | 0.78 | 0.79 | 0.48 |
| Rchi26406 | 28 | 0.76 | 0.76 | 0.45 |
| Rchi26522 | 22 | 0.62 | 0.62 | 0.41 |
| Rchi26643 | 23 | 0.87 | 0.87 | 0.68 |
| Rchi26742 | 33 | 0.88 | 0.88 | 0.52 |
| Rchi28123 | 17 | 0.8 | 0.8 | 0.64 |
| Rchi2815 | 35 | 0.9 | 0.91 | 0.59 |
| Rchi28952 | 11 | 0.69 | 0.69 | 0.74 |
| Rchi29027 | 24 | 0.89 | 0.9 | 0.72 |
| Rchi29101 | 18 | 0.66 | 0.66 | 0.5 |
| Rchi3005 | 43 | 0.94 | 0.95 | 0.67 |
| Rchi30182 | 22 | 0.68 | 0.69 | 0.45 |
| *Mean for all* | *30.1* | *0.81* | *0.82* | *0.58* |

**Supplementary Figure 1.**
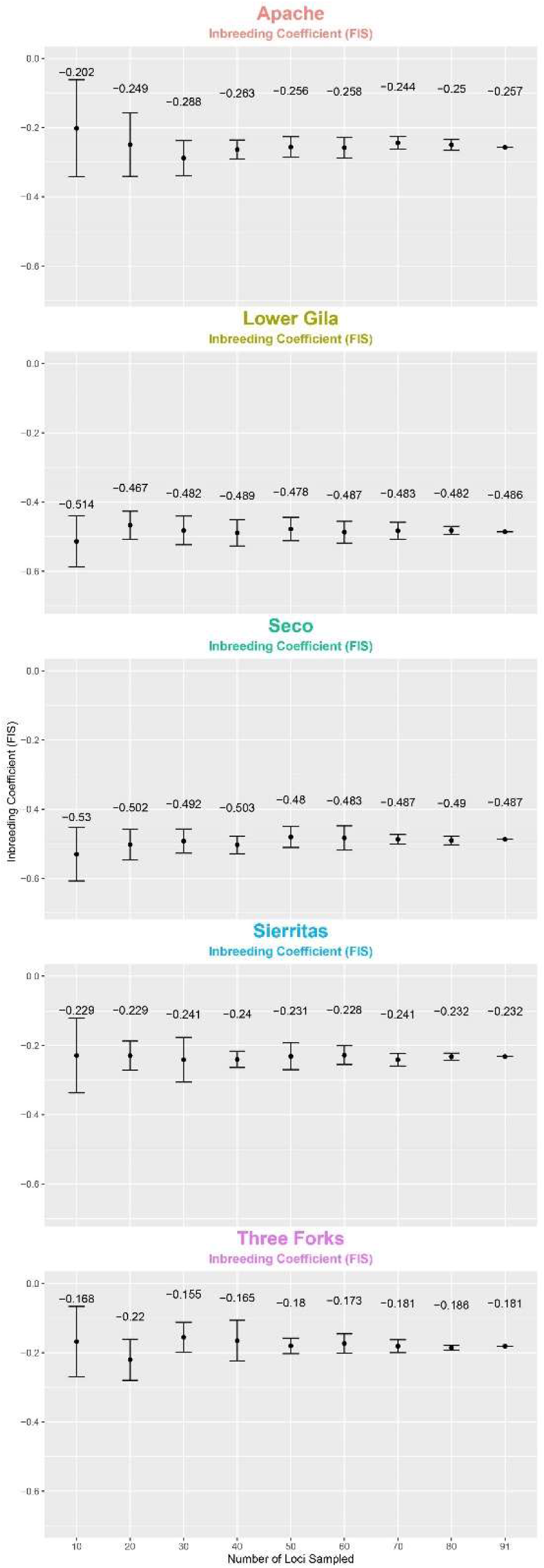
Subsampling loci show an increase in confidence for Inbreeding Coefficient (Fis) as the number of loci increase.

**Supplementary Figure 2.**
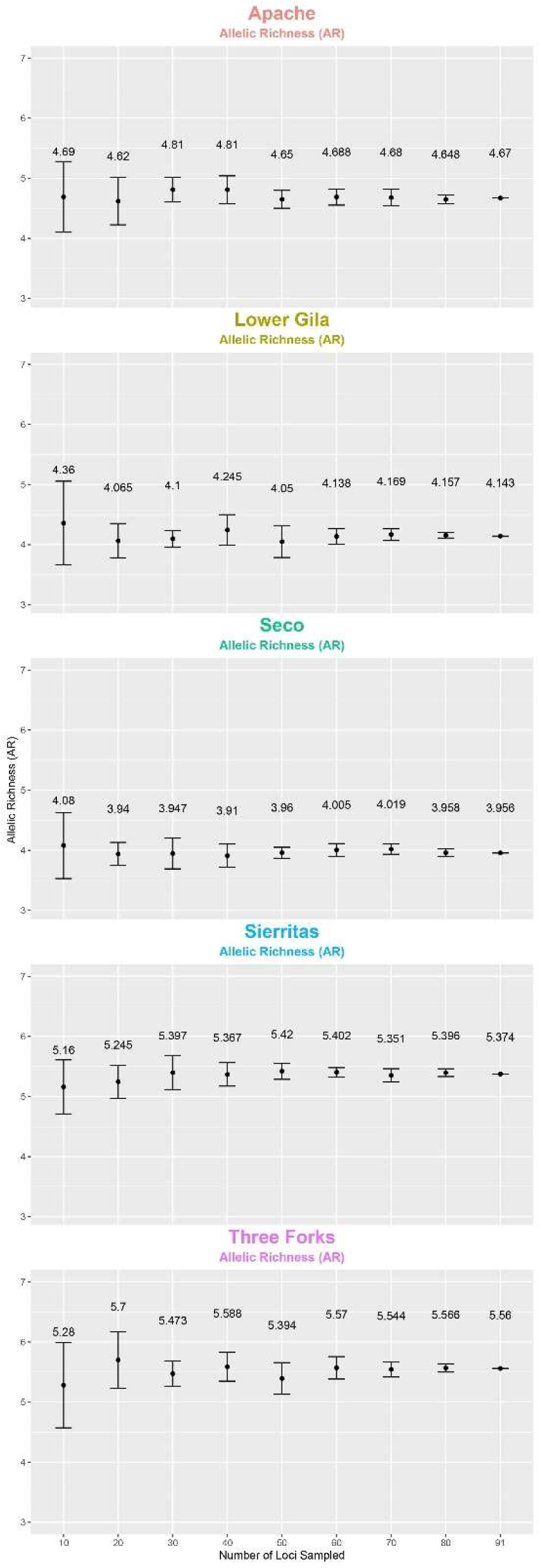
Subsampling loci show an increase in confidence for Allelic Richness (AR) as the number of loci increase.

